# Non-invasive MRI mapping of tissue-CSF water exchange reveals glymphatic fluid movement in live human cortex

**DOI:** 10.64898/2026.03.26.714533

**Authors:** Zihan Wang, Zhiyi Hu, Dengrong Jiang, Jie Song, Yifan Gou, Wen Shi, Jiani Wu, Cuimei Xu, Oluwateniola Akinwale, Kaisha Hazel, George Pottanat, Yulin Ge, Thomas Wisniewski, Vivek Yedavalli, Haris I. Sair, M. Haroon Burhanullah, Paul Rosenberg, Hanzhang Lu

## Abstract

Efficient metabolic waste clearance, via the postulated glymphatic system, is essential for neural homeostasis. However, direct visualization of tissue-cerebrospinal fluid (CSF) exchange remains limited, leading to ongoing debate in the neuroscientific field. The present work revealed evidence of tissue-CSF water exchange in the live human cortex, by employing a novel MRI technique demonstrating the flux of water molecules across the perivascular interface. We observed robust water exchange inside the cortical ribbon, which was more prominent than white matter and deep brain tissue. We validated that the signal originates from CSF and is independent of cerebral perfusion. Water exchange between tissue and CSF declined with age. Furthermore, we demonstrated for the first time that tissue-CSF exchange was impaired in Alzheimer’s disease (AD), in particular in regions where the perivascular space is clogged by anti-amyloid immunotherapy.

## Introduction

Efficient clearance of metabolic waste is essential for maintaining neural homeostasis and preventing the accumulation of neurotoxic solutes in the brain^1, 2^. Traditionally, the central nervous system was thought to lack a conventional lymphatic drainage system, but the discovery of the glymphatic system has transformed our understanding of brain fluid dynamics and waste removal^3, 4, 5^. Evidence suggests that a perivascular network facilitates the exchange of cerebrospinal fluid (CSF) and interstitial fluid (ISF)^6, 7^, promoting the clearance of soluble proteins such as amyloid-β, phosphorylated tau, and alpha synuclein implicated in the pathogenesis of neurodegenerative diseases including Alzheimer’s, Parkinson’s, and other neurodegenerative diseases^8, 9, 10, 11, 12, 13, 14^. While there is a growing recognition of the glymphatic system as a key component of brain health, there is a scarcity of direct evidence in live humans to show the fluid exchange in the cerebral cortex, which has led to some debate in the field^15, 16, 17^. Therefore, developing clinically feasible tools to assess glymphatic function and concretely demonstrating the presence of fluid exchange at the tissue-CSF interface have become pressing research priorities.

Magnetic resonance imaging (MRI) is a powerful tool to visualize human brain anatomy and function. However, it has not been able to image the glymphatic function in clinical practice. Some studies employed contrast-enhanced MRI in rodents, in which contrast agents are introduced into the CSF via intracranial^9, 18, 19, 20^ injection to trace fluid movement over time. While powerful, this approach is invasive and limited by regulatory constraints and potential neurotoxicity.

In this study, we developed a novel MRI technique for glymphatic imaging. Critically, the technique is completely non-invasive with a scan time feasible for clinical patients. Importantly, our technique employs a “remote labeling” scheme, which minimizes off-target labeling and generates “zero-background” in the image contrast. This is a distinct advantage of this technique compared to “local labeling” approaches, in which either CSF or parenchyma are labeled based on their MR properties such as T1, T2, or magnetization transfer but often suffers from unintended labeling and non-specific background signals^21, 22, 23^.

The mechanism of our technique is illustrated in Fig. 1. We use radiofrequency pulses to non-invasively label the water in the major arteries around the neck region. We purposedly ensured a sufficient spatial distance between the labeling site and the imaging site, so that there are minimal non-specific signals in the final image. These labeled water molecules are brought into the tissue space via brain perfusion, where they serve as a non-invasive “tracer” to allow the monitoring of the tissue-CSF exchange. If there is no exchange, then all tracer molecules will remain in the tissue. On the other hand, if exchange takes place, a portion of the tracer will enter the CSF space. Since CSF and tissue have characteristically different T2 MR properties^24, 25^, they can be differentially measured using tailored pulse sequences.

**Fig. 1:**
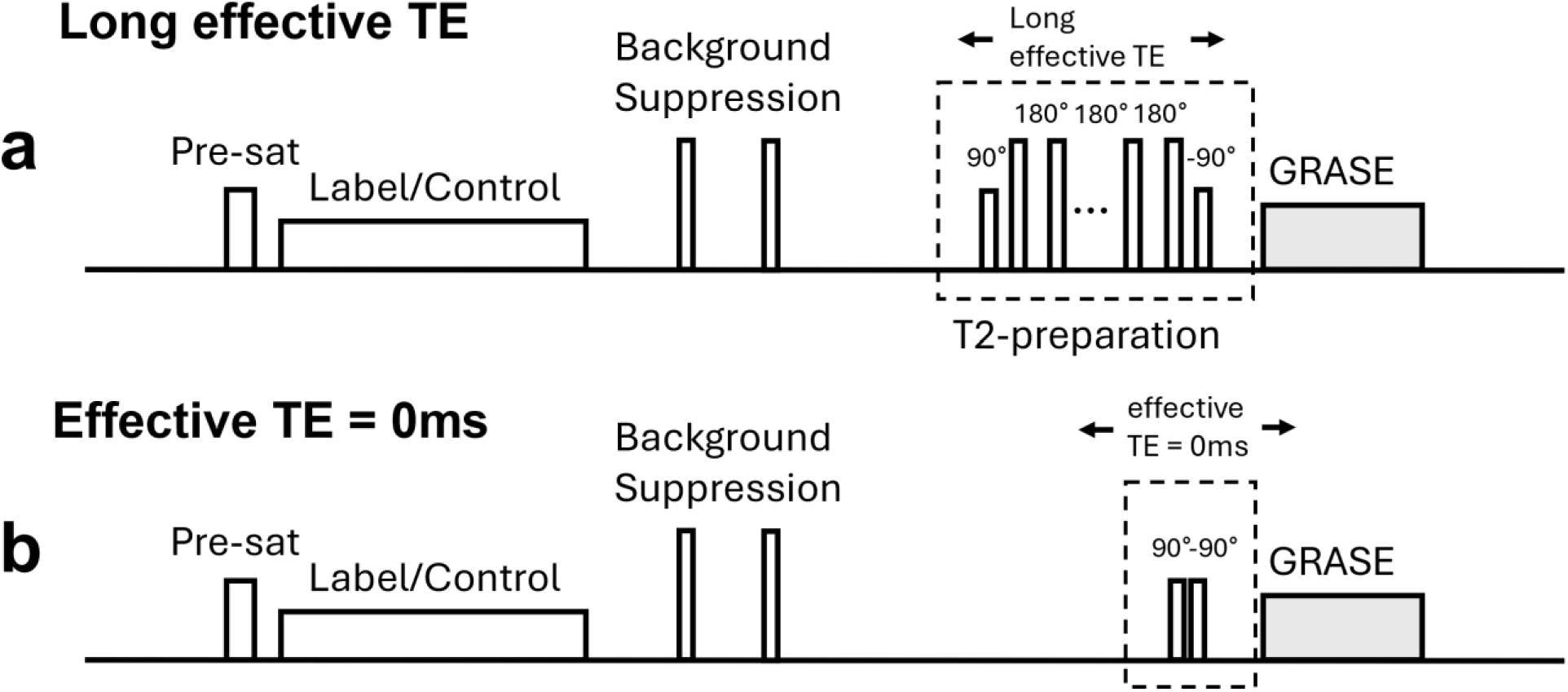
Schematic diagram of the T2-prepared long-effective-TE ASL MRI pulse sequence. **a Long effective TE sequence**. The sequence utilizes a Pseudo-Continuous Arterial Spin Labeling labeling module followed by two background suppression pulses. A non-slice-selective T2-preparation module, employing composite pulses in an MLEV pattern, is placed before the GRASE acquisition to selectively preserve signals from ASL-labeled water in the long-T2 perivascular CSF space while dissipating short-T2 tissue signals. **b Reference sequence**. A reference sequence with eTE = 0ms is performed using a composite tip-down-and-tip-up pulse to provide a reference perfusion signal, which originates from both CSF and tissue space.

We demonstrated its performance in human participants and revealed distinct spatial patterns supporting the presence of fluid exchange advanced by the glymphatic hypothesis. The robustness of the techniques was assessed in test-retest examinations. We validated its signal mechanism under a biophysical model. We further employed a physiological challenge of hypercapnia to elucidate that the glymphatic exchange rate is independent of the individual’s brain perfusion level. We also investigated age-related changes in CSF exchange. Finally, we demonstrated preliminary evidence of this technique’s clinical utility in two AD patients receiving anti-amyloid immunotherapy.

## Results

### Long-TE ASL perfusion image reveals water exchange between tissue and perivascular space

We employed a long-echo-time (long-TE) pseudo-continuous arterial spin labeling (ASL) MRI sequence by applying a T2-preparation module before the image acquisition. (Fig. 1a). This sequence aims to selectively image ASL-labeled water molecules that have exchanged into the perivascular cerebrospinal fluid (CSF) space. This is because CSF T2 (∼2,000ms) is more than one order longer than the tissue T2 (∼100ms). Thus, at an appropriate TE, the ASL signals in the tissue have completely dissipated, while those in the perivascular space are largely preserved (Extended Data Fig. 1).

The sequence was first applied to eight healthy participants (22.9 ± 2.1 years, 3 males, 5 females). We tested three eTE values of 480, 720, and 1080ms (Fig. 2a). At these eTEs, we anticipate that the tissue signal will have decayed to 0.483%, 0.034%, 0.001% of the eTE0 signal, respectively (Extended Data Fig. 1), whereas the CSF signal will be 77.5%, 68.3% and 56.4% of its eTE0 signal. For comparison, a reference ASL sequence with an eTE = 0ms (Fig. 1b) was also acquired. As shown in Fig 2a., at eTE = 0ms, the ASL maps reflect brain perfusion. At the long eTEs, it can be initially appreciated that the amplitude of the ASL signal has reduced to approximately one order lower than the original ASL signal. Nonetheless, CSF ASL signals can be reliably measured in the brain, suggesting that a certain fraction of the ASL spins have indeed exchanged into the CSF-filled perivascular space. Further visual inspection of the spatial pattern of long-TE ASL map relative to the eTE0 ASL map suggested that long-TE ASL signal is diminished in deep gray matter, e.g., caudate, putamen, and thalamus, when compared to cortical gray matter (Extended Data Fig. 2). On the other hand, long-TE ASL signal was elevated in interhemispheric fissure, sylvian fissure, and cisterns. We found that the white matter contains minimal long-TE signals.

**Fig. 2:**
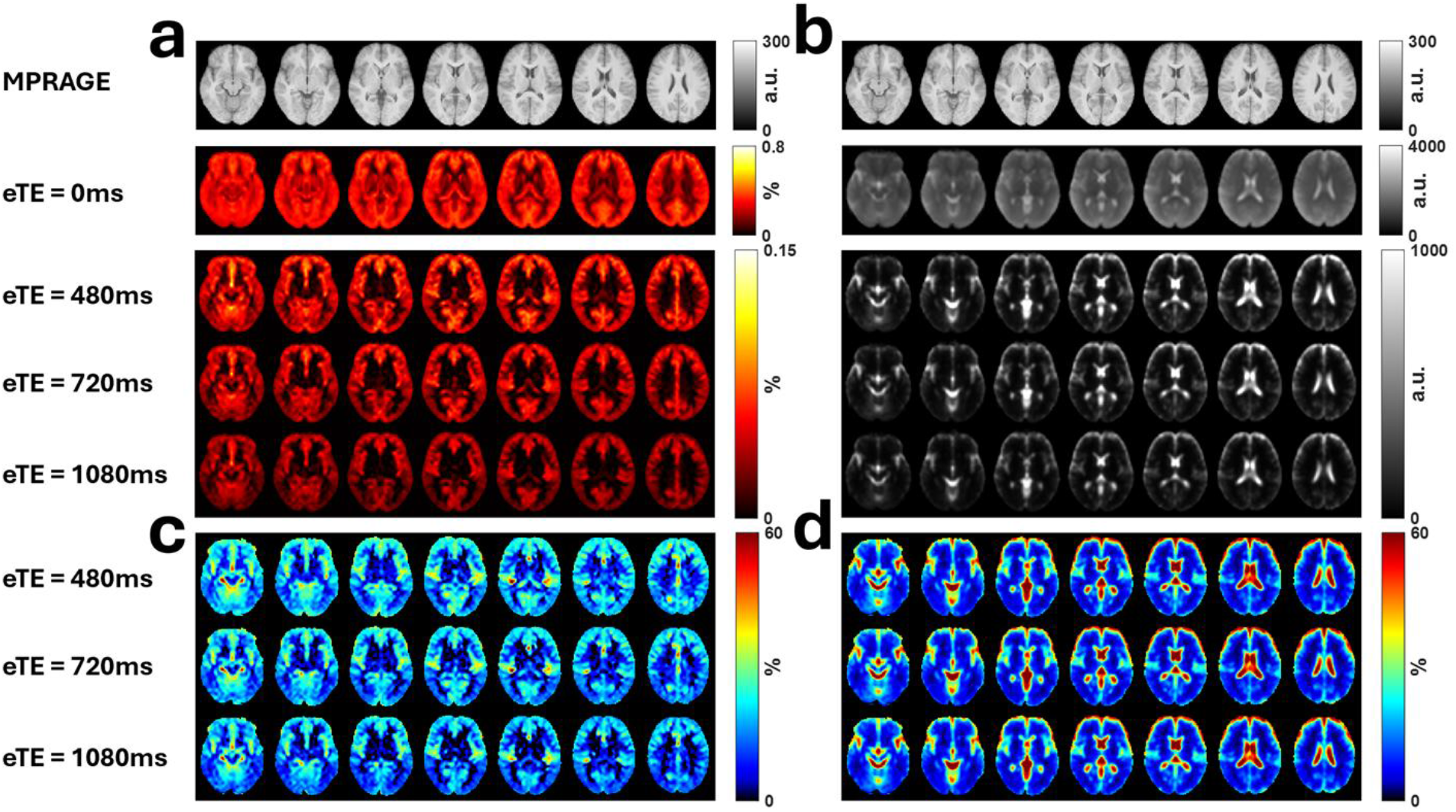
Quantitative mapping of water exchange between tissue and CSF. **a Arterial spin labeling (ASL) maps with different eTE**. The top row shows MPRAGE structural scans for anatomical reference. At eTE = 0ms, the ASL maps reflect standard brain perfusion. At long eTE (480–1080ms), the signal represents ASL-labeled spins that have exchanged into the CSF-filled perivascular space. Long-TE signal is prominent in the cortical gray matter and penetrates deep into the cortical ribbon. **b Corresponding M0 images**. At eTE = 0ms, images show spin-density contrast, while long-eTE M0 images show regions rich in CSF, most prominently subarachnoid spaces and ventricles. **c ASL CSF signal fraction**. Group-averaged maps show the ratio of long-TE ASL to eTE = 0ms signal, corrected for expected T2 decay in CSF (T2CSF = 1885.5ms). This index accounts for perfusion variations in the brain and allows the concentration on water exchange. **d M0 CSF fraction**. Maps indicate the anatomical distribution of CSF independent of the presence of exchange. The visual disparity between **c** and **d** highlights that the ASL CSF fraction specifically captures tissue-PVS exchange whereas the M0 CSF fraction identifies anatomically rich CSF regions.

Corresponding M0 images are shown in Fig. 2b. At eTE = 0ms, M0 images displayed a spin-density image contrast, while at longer eTEs, tissue signals were largely attenuated and only CSF-rich regions such as ventricles and subarachnoid spaces remained visible. Note that, in the long-TE M0 image, CSF signal is only seen at the surface of the brain. In contrast, in the long-TE ASL image, CSF signal penetrates considerably deeper into the cortical ribbon reaching the gray-matter junction. This suggests that long-TE ASL signal originates from the exchangeable perivascular space which is part of the gray matter, whereas long-TE M0 signal mainly originates from non-exchangeable subarachnoid space locating on the surface of the cortex.

### The development of a quantitative index for tissue-perivascular exchange

Next, we proposed an index to quantify the tissue-perivascular exchange, referred to as ASL CSF signal fraction. Specifically, the ASL CSF signal fraction was calculated by the ratio between long-TE ASL signal and eTE0 ASL signal, with corrections of expected T2 decay in CSF. This index allows the separation of the effects of exchange from perfusion. Also note that we only used one long-TE data to compute this index, rather than using multiple long-TE data, to ensure an affordable scan time and the clinical translation potential of the technique. Fig. 2c shows the group-averaged ASL CSF fraction map. Fig. 2d shows a M0 CSF fraction map computed similarly but used the M0 images, which indicates the anatomic distribution of CSF in the brain irrespective of their exchange property. It can be seen that the ASL CSF fraction and M0 CSF fraction maps are visually disparate. The ASL CSF fraction has a strong signal in the cortical gray matter, where the glymphatic exchange is thought to be most prominent. The ventricles, deep gray matter, and white matter were found to have low ASL CSF fractions.

The ASL CSF fraction maps obtained with different eTE values are largely similar under visual inspection (Fig. 2c). Fig. 3a shows quantitative values of ASL CSF fraction in different ROIs (Extended Data Fig. 3). For cortical gray matter, there was a slight decline with increasing eTE, likely reflecting incomplete tissue signal suppression at shorter T2-preparations. Test-retest reproducibility assessment revealed a coefficient-of-variation (COV) of 6.8% ± 3.3%, 6.5% ± 4.1%, and 9.0% ± 4.6% for the three eTEs. Considering a tradeoff between signal accuracy and reproducibility, eTE = 720ms was selected for subsequent experiments. For comparison, the M0 CSF fraction values are shown in Fig. 3b. These values are characteristically different from the ASL CSF fraction values.

**Fig. 3:**
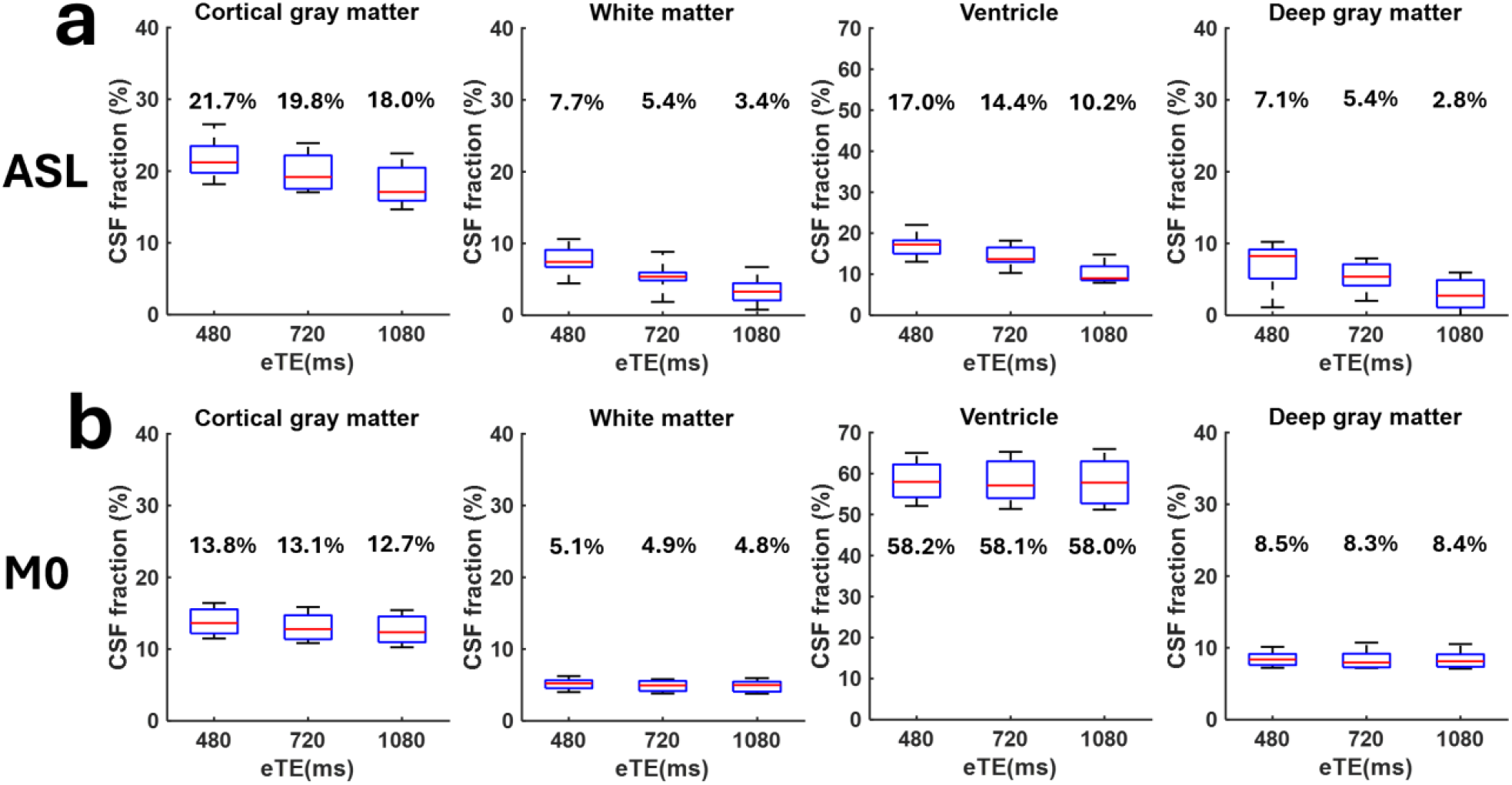
Quantitative regional values of CSF fraction. **a ASL CSF signal fraction across regions of interest (ROIs)**. Boxplots show the distribution of the ASL-based exchange index in cortical gray matter, white matter, ventricles, and deep gray matter at three eTE values. Cortical gray matter exhibits the highest exchange fraction, with a slight decline at longer eTE potentially due to reduced tissue signal contamination. **b CSF volume fraction across ROIs**. Quantitative values derived from M0 images show a characteristically different distribution compared to ASL-based maps, with the highest values located in the ventricles.

### Verification that the long-TE ASL signal is originated from the CSF space

To further verify that the long-TE ASL signal is indeed located in the CSF space, we exploited the T1 differences between CSF and tissue. We reasoned that, since the labeling effect decays with the T1 of the spin which is determined by its microenvironment, the long-TE ASL spins should decay at CSF T1 whereas the eTE0 ASL spins should decay with the tissue T1. We therefore examined how the long-TE ASL signal is dependent on post-labeling delay (PLD). Nine healthy volunteers (25.3 ± 3.4 years; 4 males, 5 females) were scanned at PLDs ranging from 2500ms to 4000ms. Group-averaged ASL difference images for both eTE0 and eTE720 are shown in Fig. 4a,b, respectively. The eTE0 images revealed a rapid intensity decrease with PLD. On the other hand, the eTE720 images were relatively stable. Quantitative values from the cortical gray matter are also shown in Fig. 4a,b. Exponential fitting resulted in the decay time-constant of eTE0 and eTE720 signals, which was found to be 1715ms and 5261ms, respectively. These values are consistent with T1 of the corresponding spin compartments, predominantly tissue compartment for eTE0 and CSF compartment for eTE720 data. These observations confirm that long-TE ASL signals reflect the amount of ASL spins that have been exchanged into the CSF-filled perivascular space.

**Fig. 4:**
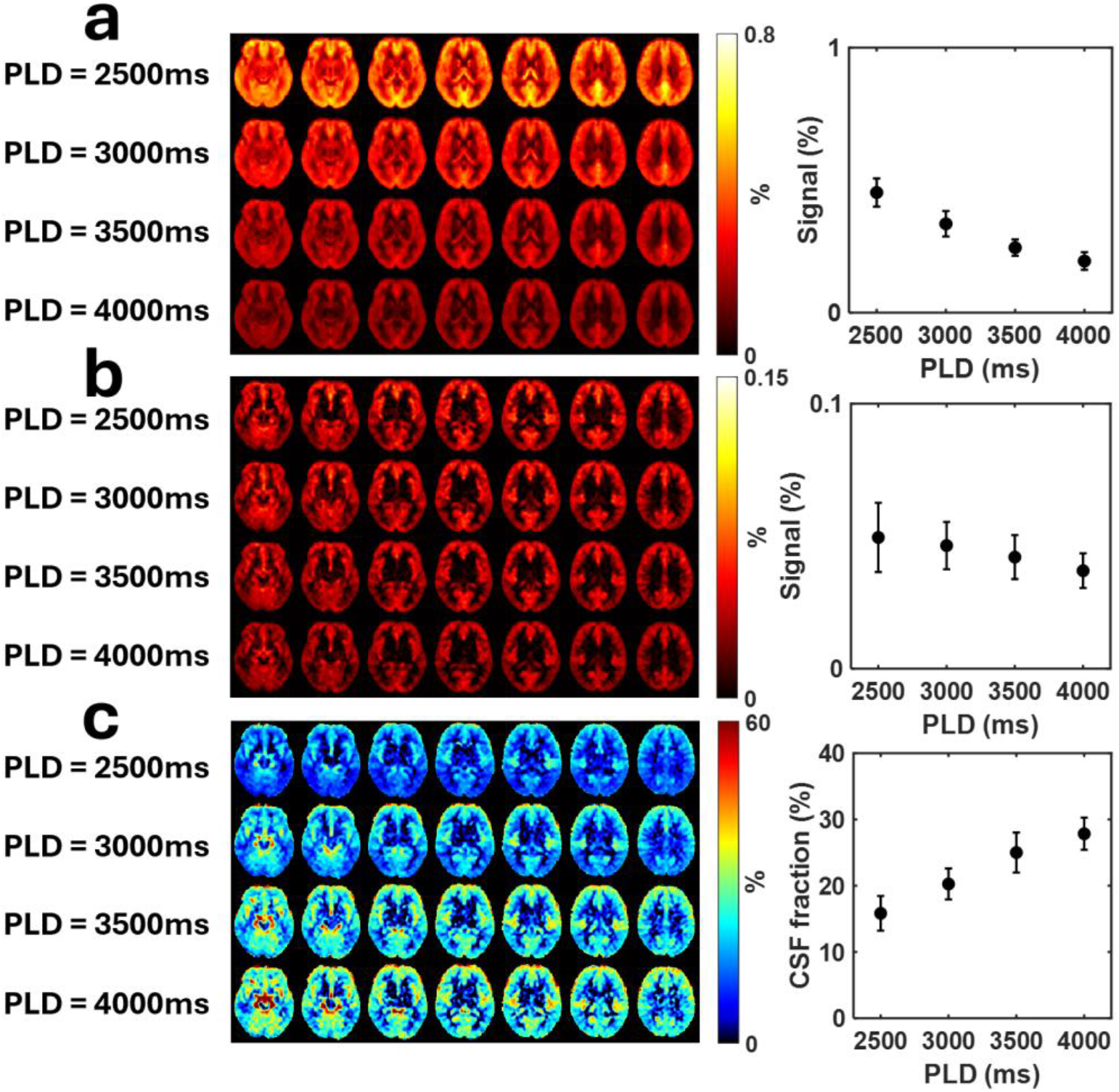
Verification of signal origin based on its T1 property. **a Post-Labeling-Delay (PLD) dependence of eTE = 0ms ASL signal**. The signal shows rapid intensity decrease with a decay time-constant of 1715ms, consistent with tissue T1. **b PLD dependence of eTE = 720ms ASL signal**. The signal remains relatively stable with a decay time-constant of 5261ms, consistent with the long T1 of the CSF compartment. **c ASL CSF fraction vs. PLD**. The fraction increases from 15.8% to 27.8% as PLD increases from 2500ms to 4000ms, reflecting the differential T1 decay rates of tissue and CSF compartments. These findings confirm the long-TE ASL signal originates from the CSF-filled perivascular space.

The ASL CSF signal fraction maps are shown in Fig. 4c. The fractions became greater at higher PLD, rising from 15.8 ± 2.6 % at 2500ms to 27.8 ± 2.4 % at 4000ms. The dependence of ASL CSF signal fraction on PLD can be expected from the T1 differences between tissue and CSF.

### Hypercapnia challenge via CO2 inhalation increases brain perfusion but does not alter tissue-perivascular exchange rate

To verify that the proposed ASL CSF signal fraction represents a physiological index distinct from conventional brain perfusion, we conducted a hypercapnia challenge in six healthy adults (23.2 ± 0.8 years; 2 males and 4 females). Hypercapnia challenge via CO2 inhalation is known to increase cerebral blood flow. It can be seen that, CO_2_ inhalation produced robust perfusion-related signal increases in both eTE0 and eTE720 ASL acquisitions (Fig. 5a,b), by 12.5 ± 8.2% (p=0.010) in eTE0 and 10.2 ± 8.0% in eTE720 (p=0.019). In contrast, the ASL CSF signal fraction remained unchanged (17.4% ± 1.1% under room air vs. 17.3% ± 3.1% during hypercapnia; p = 0.890) (Fig. 5c). This observation is consistent with the notion that the tissue-perivascular exchange rate is a property of the CSF-brain barrier and is independent of the perfusion to the brain. Cerebral perfusion level only changes the amount of “tracer” applied in the measurement, but does not affect the exchange index itself.

**Fig. 5:**
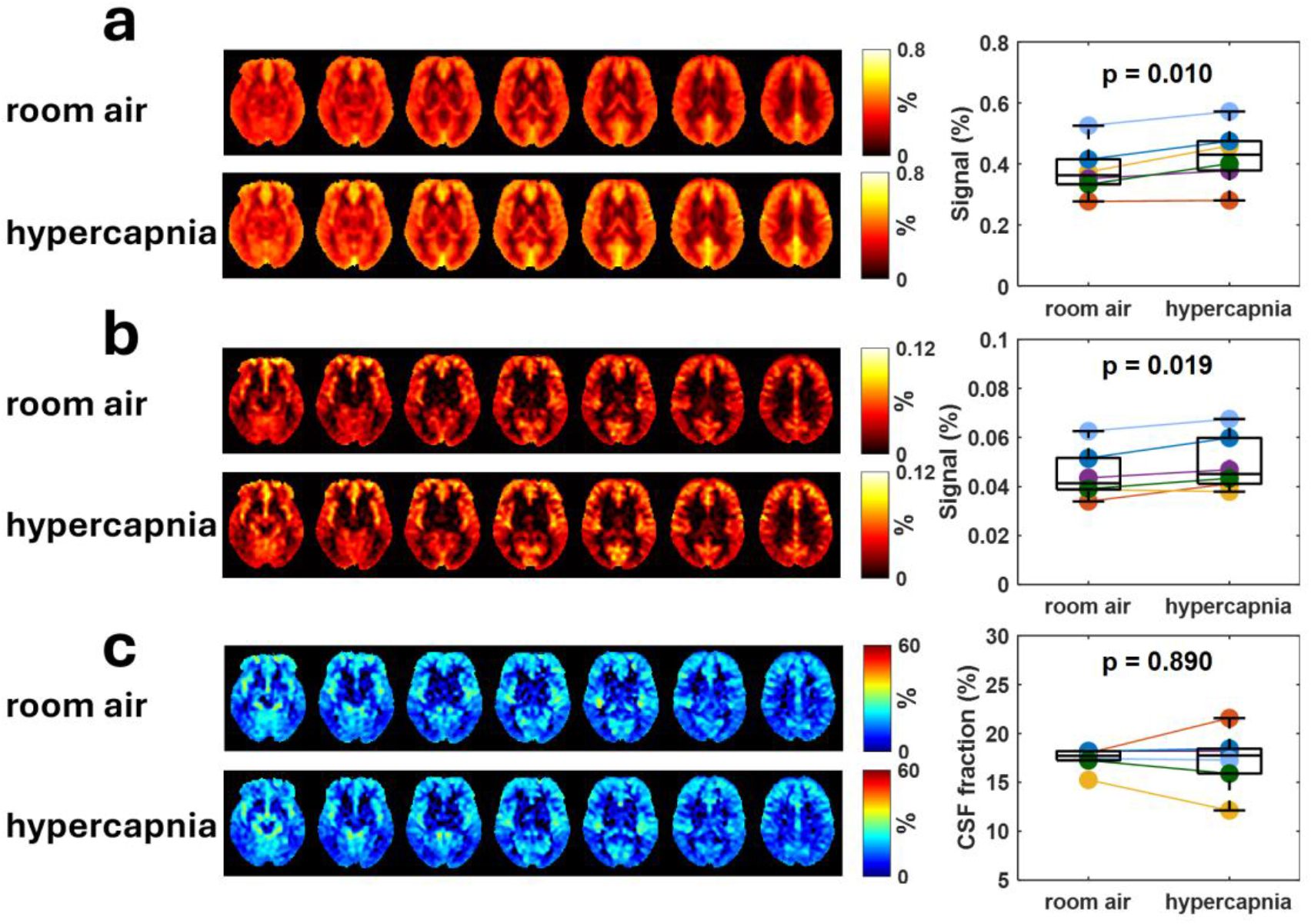
Independence of tissue-CSF exchange from cerebral perfusion. **a eTE = 0ms ASL signal**. Hypercapnia via CO_2_ inhalation produced a 12.5±8.2% (p=0.010) increase in perfusion signal. **b eTE = 720ms ASL signal**. Long-TE signal increased by 10.2±8.0% (p=0.019) during hypercapnia. **c ASL CSF signal fraction**. The exchange index remained unchanged between room air and hypercapnia (p=0.890), indicating that the exchange rate is a property of the interstitial fluid (ISF)-CSF interface independent of total brain perfusion.

### Brain aging is associated with an increase in CSF volume fraction but a reduction in water exchange between tissue and perivascular space

We investigated age-related differences in long-TE ASL MRI from a cohort of 37 healthy participants aged 21–82 years. Multiple linear regression revealed an age-related decrease in ASL CSF signal fraction (slope = –0.07 ± 0.03, p = 0.013) (Fig. 6a), suggesting that the exchange of water between interstitial tissue and perivascular space is diminished in aging. In contrast, M0 CSF fraction increased with age (slope = 0.15 ± 0.02, p < 0.001) (Fig. 6b), consistent with age-related brain volume reduction and CSF volume increase. No significant associations were observed between sex and CSF fraction in either measurement.

**Fig. 6:**
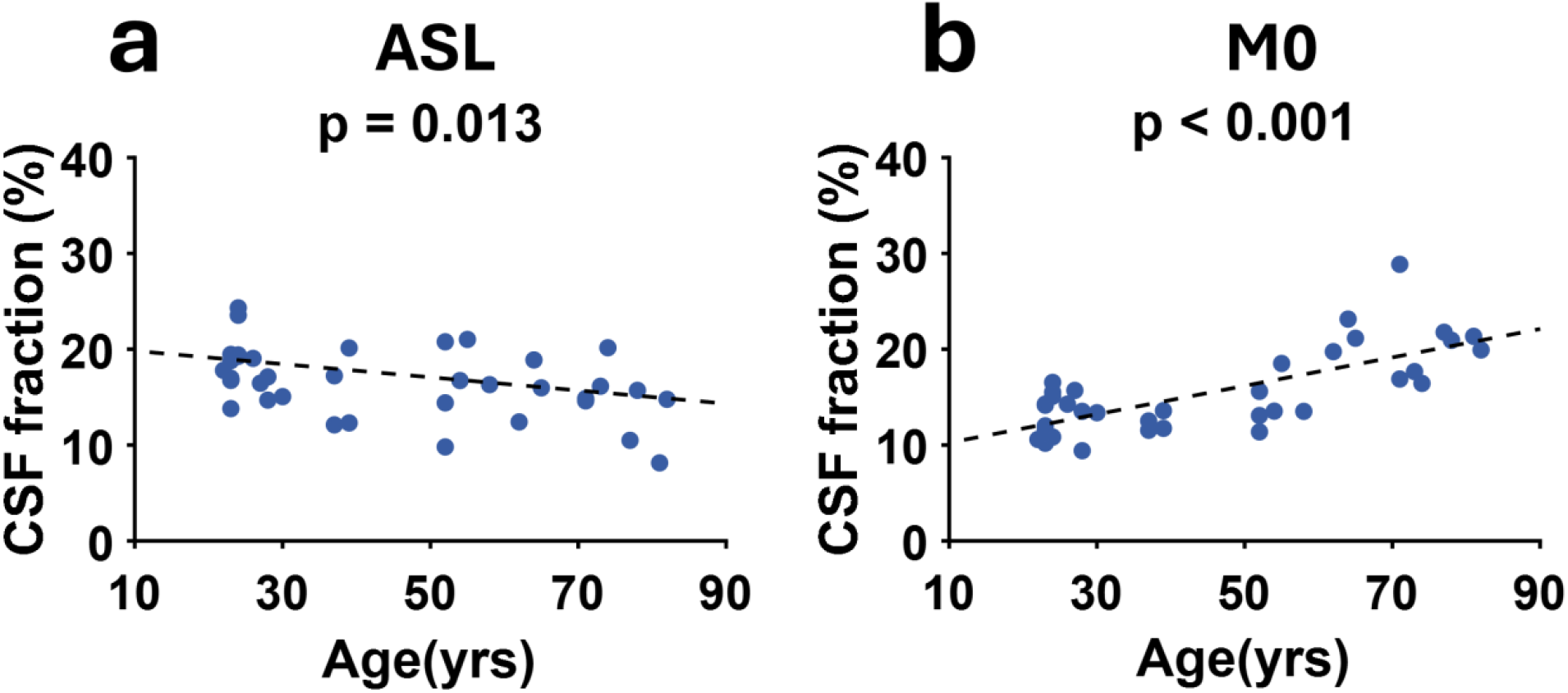
Age-related decline in glymphatic exchange. **a Age-related changes in ASL CSF fraction**.Multiple linear regression in a cohort of 37 healthy participants (21–82 years) reveals a significant decrease in the exchange index with age (slope=–0.07±0.03, p=0.013), indicating diminished water exchange between interstitial tissue and the PVS. **b Age-related changes in M0 CSF fraction**. In contrast, the M0 CSF fraction increases with age (slope=0.15±0.02, p<0.001), consistent with age-related brain volume reduction and increased total CSF volume.

### Case study: Impaired water exchange between tissue and perivascular space in patients with amyloid-related imaging abnormalities

We applied the proposed sequence to two patients who underwent anti-amyloid immunotherapy with donanemab and subsequently developed amyloid-related imaging abnormalities (ARIA). The first patient is an 83-year-old male patient who completed three infusions administered at weeks 0, 4, and 8. ARIA was developed at week 10, prompting cessation of treatment. A research MRI was performed at week 16. Imaging findings from this patient are summarized in Fig. 7 (left panel), with the treatment timeline shown in Fig. 7a. ARIA was first detected at week 10 on clinical T2-FLAIR imaging and susceptibility weighted imaging, after which further infusions were discontinued. Serial clinical T2-FLAIR and susceptibility-weighted imaging demonstrated both progressive edema (ARIA-E) and hemorrhage (ARIA-H) changes (Fig. 7b). Research MRI in week 16 revealed markedly reduced long-TE ASL signal around the ARIA-E and ARIA-H region (Fig. 7c).

**Fig. 7:**
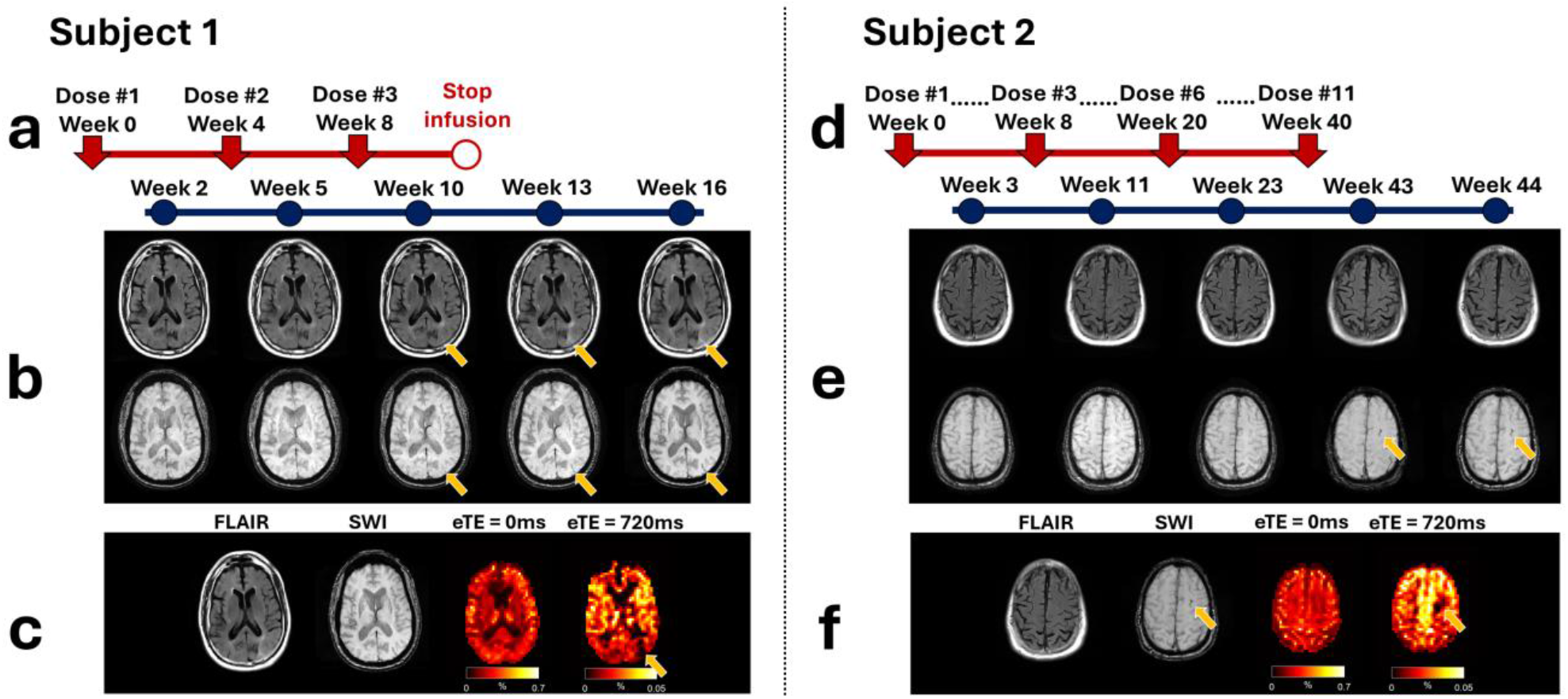
Clinical application in patients with Amyloid-Related Imaging Abnormalities (ARIA). **a–c Subject 1 (83-year-old male)**. Treatment timeline (**a**) and clinical imaging (**b**) showing edema (ARIA-E) and hemorrhage (ARIA-H) following donanemab infusions. Research MRI at week 16 (**c**) reveals markedly reduced long-TE ASL signal and diminished CSF signal fraction in the affected regions. **d–f Subject 2 (75-year-old male)**. Treatment timeline (**d**) and clinical imaging (**e**) showing ARIA-H at week 43. Research MRI at week 44 (**f**) demonstrates reduced long-TE ASL signal and diminished CSF signal fraction in ARIA-H region.

The second patient is a 75-year-old male patient who completed eleven monthly infusions. Following his eleventh infusion at week 40, a clinical MRI performed at week 43 revealed new ARIA-H abnormalities. A research scan was subsequently performed in week 44. Imaging findings from this patient are summarized in Fig. 7 (right panel), with the treatment timeline shown in Fig. 7d. ARIA-H was first detected at week 43 on susceptibility weighted imaging. Both clinical scan in week 43 and research scan on week 44 showed stable microhemorrhage and superficial siderosis (Fig. 7e). Research MRI in week 44 revealed markedly reduced long-TE ASL signal around the ARIA-H region (Fig. 7f).

Together, these observations suggest impaired water exchange at the glymphatic interface in regions affected by ARIA.

## Discussion

In this study, we demonstrate for the first time that tissue-CSF exchange, the core component supporting the glymphatic hypothesis, can be measured in a clinically practical setting with no off-target labeling. This novel technique was able to generate voxel-wise maps of the tissue-CSF exchange with high quality and reproducibility, in less than 10 minutes on a standard MRI system. We verified the signal mechanism of our technique using both a biophysical model and a physiological challenge of hypercapnia. As an initial demonstration of the clinical utility of this new technique, we applied it to older adults and two AD patients who received anti-amyloid immunotherapy and subsequently developed ARIAs. We found that the tissue-CSF exchange was diminished with aging and its further impairment in AD was spatially co-localized to regions with an amyloid-clogged tissue-CSF interface.

Prior to the proposal of a glymphatic hypothesis, it was thought that the primary location that CSF can exchange with the brain is the choroid plexus^26, 27, 28^. The ASL CSF fraction map indeed confirmed the presence of an exchange signal in the choroid plexus. More recently, the glymphatic system emerged as an exciting topic in understanding the mechanisms underlying neurodegenerative diseases, which hypothesizes that there exists a vast exchange of water between the brain parenchyma and CSF in the perivascular space. Furthermore, such water movements bring along with them metabolic waste products, such as beta-amyloid; in health, this leads to effective clearance of these toxic peptides. Our observation of rich ASL CSF signals in the cortical ribbon, provides strong support for the presence of water exchange beyond the choroid plexus and aligns with the glymphatic hypothesis. The spatial pattern of the ASL CSF fraction is particularly striking when compared to the M0 CSF fraction (i.e., the anatomic distribution of CSF in the brain), which primarily revealed bright CSF signal on the surface of the cortex where subarachnoid space is. The ASL CSF fraction clearly penetrates much deeper into the cortex, suggesting that, while the subarachnoid space contains substantial CSF, this is not in exchange with brain tissue. On the other hand, the perivascular space around the penetrating arteries and veins is a smaller CSF-filled space but it is in rapid exchange with brain tissue water.

Clinically, there are no existing techniques to noninvasively measure glymphatic exchange in live humans. Contrast based methods in humans, including intracranial or intrathecal injections, are highly invasive and not applicable to routine clinical practice. The proposed technique is based on a similar labeling principle as the ASL technique, and has the advantage of being high specific to tissue-CSF exchange. Compared to other techniques such as T2-labeling and MT-labeling, our technique has no contamination by non-specific background signals due to unintended labeling. Furthermore, unlike a previous study which suffered from low signal-to-noise ratio (SNR) and require extra-long scan time (50 minutes)^29, 30^, our technique is based a new noise-attenuation design referred to as CSF background suppression and also employs a cutting-edge T2-preparation 3D GRASE acquisition. These technical advances collectively enhanced the sensitivity and efficiency of our technique. As a result, we are able to obtain a high-quality map at long TE even at a signal level 0.1% of the equilibrium magnetization. Another major SNR-boosting approach we employed was a single-shot, as opposed to segmented, 3D acquisition. In conventional ASL sequence, single-shot 3D acquisition was not desired because tissue T2 is relatively short and the long echo-train in single-shot acquisition will result in considerable signal decay. Therefore, segmented acquisition was used in previous studies^29^, which increased the scan time 8 fold. We leveraged the uniquely long T2 of CSF and showed that single-echo acquisition can still preserve the signal and spatial acuity. In summary, this highly sensitive technique is clinically suitable for repeated scans, without any radiation or contrast agent, which could be part of large-cohort studies and longitudinal studies.

Given the complexity of the glymphatic system and its poorly understood physiological underpinnings, it is important to validate the mechanisms of our technique. In this report, we conducted two studies to verify the source of our signals. In one study, we reasoned that, if our measured signal indeed originates from the CSF space, its time-dependent decay will follow the corresponding micro-environment, i.e., T1 characteristic of CSF. Supporting our hypothesis, our delay-dependent study revealed an intrinsic T1 of 5261ms for the long-TE ASL signal, which was drastically longer than the T1 (1715ms) of standard ASL signal. This study verified that the source of the signal of our long-TE ASL sequence is the labeled spins located in the CSF space. We also wanted to confirm that our hypothesized signal mechanism is not contaminated by other physiological parameters. Since our pulse sequence is modified from the ASL sequence, one important question is whether the signal we obtained is just another form of perfusion signal. To rule out this possibility, we employed a hypercapnia challenge which is known to increase the perfusion to the brain. We postulate that, since our technique aims to measure water exchange rate, which is a property of the tissue-CSF interface independent of perfusion, our signal should not be altered by the hypercapnia challenge. Our results suggested that, despite a perfusion increase, the ASL CSF fraction was not altered by hypercapnia, providing further support for the validity of the ASL CSF fraction proposed in this report.

One potential confounding factor is that the labeled arterial water may enter the perivascular space directly from the blood vessels, rather than entering the tissue then reaching the perivascular space via ISF-CSF exchange. While this is a plausible possibility, data from our multi-PLD experiments suggest that the majority of the labeled water in the perivascular space comes from the ISF. Specifically, the long-TE ASL signal in the perivascular space decayed slowly with PLD (within a PLD range of 2500 to 4000ms), with a decay time constant longer than the T1 of CSF. If the majority of the labeled spins in the perivascular space had come from the vessel, the signal should decay more rapidly, due to a combined effect of T1 relaxation and loss of labeled spins (from CSF to ISF). In contrast, our observation of a long decay constant suggests that the perivascular space gained labeled spins (from ISF to CSF) in this PLD range. Additionally, neuroanatomy knowledge taught us that the arterial wall is largely impermeable to water^31, 32^ and capillaries have minimal perivascular space compared with arteries or veins^10, 33^.

Therefore, most labeled spins in the perivascular space are expected to arrive from the interstitial space.

Studies of this technique in the context of aging and disease can shed additional light on the scientific utility of the proposed technique. There is growing evidence that glymphatic function is impaired with age, which may lead to poor clearance of metabolic waste and may be a mechanism underlying some neurodegenerative diseases including Alzheimer’s and Parkinsons’. This has been demonstrated in rodent models but human evidence of glymphatic dysfunction has been limited ^23, 34^. The present study observed an age-related decrease in ASL CSF fraction, providing important human evidence on a decline in waste clearance capacity in older adults.

Note that reduced ASL CSF fraction with age is in contrast with an increase in M0 CSF fraction, which is attributed to anatomic change, i.e., brain atrophy, with age. Therefore, the proposed technique provides critical insights on the brain beyond anatomic information.

Anti-amyloid immunotherapy has been one of the most exciting advances in the AD field in the past decade; representing the first FDA approved disease-modifying treatment^35^. However, a prominent side-effect is the amyloid-related imaging abnormalities, which can be fatal ^36, 37^. Therefore, it is of major clinical significance to understand its pathophysiology and identify potential biomarkers for its prediction. In the initial testing of our proposed technique in two AD patients, we observed markedly reduced signal in regions corresponding to ARIA. These observations provide preliminary evidence that tissue-CSF exchange may be impaired in ARIA, possibly due to the clogging of the ISF-CSF interface by amyloid peptides from dissolving plaques, being cleared by the immunotherapy, or due to the blockage of downstream perivascular space disrupting the CSF outflow. Therefore, our technique may not only be useful in understanding ARIAs but also for understanding the natural history of AD and treatment response to anti-amyloid immunotherapies. There may be natural variation in glymphatic clearance which could explain variation in outcomes such as rate of longitudinal decline.

Additionally, this technique could serve as a surrogate marker to aid in development of AD therapies that modify (presumably, increase) glymphatic clearance.

There are several limitations to this study. First, the current implementation does not achieve whole-brain coverage. Future improvements could extend the echo train length for enhancing the spatial coverage. Second, the proposed technique has not been compared with other methods for validation. Furthermore, although preliminary results demonstrate the sensitivity of the technique in AD, further studies in larger patient cohorts are needed. Additionally, there are some regional heterogeneities in the ASL CSF signal fraction map, e.g., the lower lateral temporal signals, that may be attributed to confounding factors such as the bolus arrival time. Finally, the present technique measures water exchange, but not the actual clearance of metabolic waste. Thus, it remains to be shown whether water and metabolites are directly related in terms of their motion.

In conclusion, we measured water motion in the live human brain, using a non-invasive MRI technique. We observed pronounced water exchange between brain tissue and CSF at the perivascular interface, providing a strong support for the convex fluid flow that is the basis of the glymphatic hypothesis. We showed that the exchange is most active in the cortical ribbon, rather than in CSF-abundant areas such as the ventricles or the subarachnoid space, suggesting a disparity between anatomically and functionally active CSF regions. Using this method, we identify a significant age-related decline in water exchange and also demonstrate initial evidence of its sensitivity to AD. Together, these findings establish a quantitative, completely non-invasive technique for human glymphatic imaging, with a strong potential for numerous translational and clinical applications.

## Methods

### General experiment design

The study protocols were approved by the Institutional Review Board of the Johns Hopkins University. Written informed consent was obtained from each participant prior to the study procedures. MRI data were acquired on a 3T Siemens Prisma system (Siemens Healthineers, Erlangen, Germany) using a body coil for transmission and a 32-channel head coil for receiving. A total of 56 participants were enrolled.

### MRI pulse sequence

The T2-prepared ASL sequence (Figure 1) is comprised of Pseudo-Continuous Arterial Spin Labeling (PCASL) module followed by two background suppression pulses. To maximumly suppress the contamination from the non-labeled CSF signal, an enhanced background suppression scheme was used and the data were stored in complex format allowing the estimation of the sign of the magnetization^38^. A T2-preparation module^39, 40^ with an effective TE (eTE) was placed before a gradient and spin echo (GRASE) acquisition to separate tissue and CSF signal contributions. The T2-prepation used non-slice-selective RF pulses to ensure that module is insensitive to flow. To minimize the imperfection of T2 preparation pulses, composite pulses were used (i.e., 90°_x_180°_y_90°_x_ for the 180° pulses) and the signs of the 180° pulses were arranged in a MLEV pattern (e.g., 1 1 -1 -1). As a reference, an equivalent sequence but using eTE = 0ms was also acquired. To achieve eTE = 0ms, a composite tip-down-and-tip-up pulse was placed before GRASE acquisition. For comparison, a M0 sequence without labeling module and background suppression pulses were acquired, with a T2-preperation module matching the ASL sequence and a TR of 10,000ms.

### Theory and MRI signal mechanism

The T2 of CSF is substantially longer than that of brain tissue, a property that can be leveraged to isolate CSF contributions from mixed MRI signals. At an eTE = 0ms, the acquired signal consists of both tissue and CSF components, expressed as:

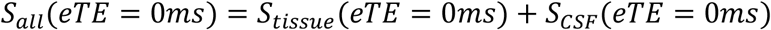

When a sufficiently long T2-preparation module is applied, the tissue component undergoes nearly complete T2 decay, whereas the CSF component, due to its prolonged T2, is largely preserved. Under these conditions, the total measured signal predominantly reflects the CSF contribution:

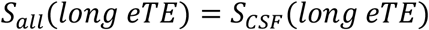

We further estimated an index of CSF fraction in ASL MRI. To account for the T2 relaxation of CSF, we corrected the measured CSF signal using an assumed T2 of 1885.5ms^24^, based on prior literature values at 3T. This correction effectively estimates the CSF signal as it would appear at eTE = 0ms. The CSF signal fraction can be calculated by dividing the corrected CSF signal by total signal at eTE0:

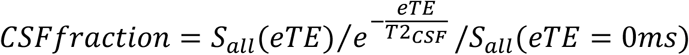

### Study I

Eight healthy participants (22.9 ± 2.1 years, 3 males, 5 females) were enrolled. Each participant underwent T2-prepared ASL sequence at four different eTEs: 0ms (reference), 480ms, 720ms, and 1080ms. Each eTE was scanned twice to allow the evaluation of test-retest reproducibility. The other imaging parameters were: labeling duration=3000ms, Tau_CPMG_ = 15ms. FOV=216×216×80mm^3^, voxel size=4×4×4mm^3^, slice oversampling = 20%, TE/TR=31.2/6830ms, excitation flip angle = 90°, refocusing flip angle = 120°, echo train length=748.8ms, number of averages=20, scan time=4min47sec for each sequence.

For anatomic reference, a T1-weighted image was acquired with the following parameters: FOV=256×224×80mm^3^,voxelsize=1×1×1mm^3^, TE/TR=4.18/2100ms, inversion time = 1100ms, flip angle = 12°, scan time=4min20sec for each sequence.

Raw ASL data were processed offline using in-house MATLAB (Mathworks, Natick, MA) scripts. ASL data were preprocessed using Statistical Parametric Mapping Version 12 (SPM12, University College London, London, UK). The control and labeled images were realigned to the first dynamic for motion correction, and image scrubbing was performed similar to fMRI data process strategies^41^. MRI signals were initially stored as complex numbers. The complex control-minus-label vectors were projected on to the direction of the M0 signal vector to obtain a signed scalar^38^.

Quantitative analysis was performed on four regions of interest (ROIs): cortical gray matter, white matter, deep gray matter, and ventricle. To obtain ROIs, tissue probability maps were generated from the M0 image acquired at eTE = 0ms using SPM12. The ROIs were then defined semi-automatically by thresholding on the tissue probability maps in combination with manual delineation (Extended Data Fig. 3).

### Study II

Nine healthy participants (25.3 ± 3.4 years, 4 males, 5 females) were scanned at multiple PLDs to examine the dependence of long-TE ASL signals on spin relaxation properties. Long-TE ASL scans were acquired at PLDs of 2500ms, 3000ms, 3500ms, and 4000ms. Both eTE = 0ms and eTE = 720ms (selected based on reproducibility analysis) conditions were obtained. M0 images with corresponding eTEs and anatomical images were also acquired. The processing methods were identical to those of Study 1.

### Study III

Six healthy participants in total (23.2 ± 0.8, 2males, 4 females), were recruited separately from Study I and Study II to investigate the effect of hypercapnia to tissue-perivascular water exchange. The long-TE ASL sequence with eTE = 720ms and the reference eTE = 0ms sequence were acquired in all participants under a normocapnia (breathing room-air) and hypercapnia (breathing CO_2_ enriched gas). The CO_2_ gas mixture contains the following contents: 4% CO_2_, 21% O_2_, and 75% N_2_.

### Study IV

Thirty-seven healthy participants in total, aged 21–82 years (45.1 ± 21.2, 17males, 20 females), were recruited separately from Study I, Study II, and Study III to investigate age-related changes in tissue-perivascular water exchange. The long-TE ASL sequence with eTE = 720ms and the reference eTE = 0ms sequence were acquired in all participants. M0 images with were also acquired.

### Study V

Two Alzheimer’s patients receiving anti-amyloid antibody therapy were included in this study.

The first patient was an 83-year-old male with Alzheimer’s disease receiving anti-amyloid therapy with donanemab, who completed three infusions administered at weeks 0, 4, and 8. At week 10, clinical MRI revealed the development of amyloid-related imaging abnormalities (ARIA). Specifically, sulcal effusion (ARIA-E) was observed in the left occipital region, along with superficial siderosis (ARIA-H) in the left occipital lobe, corresponding to areas of abnormal signal on FLAIR images. Research MRI scan was subsequently performed in week 16. At week 16, persistent edema was present in the left occipital and periventricular white matter, as well as in the parieto-occipital sulcal region, with increased effusion involving the left parieto-occipital sulcus and adjacent periventricular white matter. Additionally, an increased number of microhemorrhages were detected in the left occipital lobe surrounding the region of superficial siderosis initially observed at week 10.

The second patient was a 75-year-old male with Alzheimer’s disease undergoing anti-amyloid therapy with donanemab, who completed a total of eleven infusions. Following the eleventh infusion at week 40, a clinical MRI performed at week 43 revealed new ARIA-H abnormalities.

Specifically, a new microhemorrhage was identified as a punctate susceptibility focus within the white matter of the right frontal lobe, and new superficial siderosis was detected as a focal linear susceptibility signal along the left superior frontal sulcus. A research MRI scan was subsequently acquired at week 44, which demonstrated stable findings compared with week 43, including the right frontal lobe microhemorrhage and superficial hemosiderosis in the left superior frontal sulcus.

For research scans, the proposed long-TE ASL sequence (eTE = 720ms) and a reference sequence with eTE = 0ms were acquired. Corresponding M0 images were also obtained.

### Statistical analysis

Coefficient of variation (COV) of CSF fraction between two repeated scans was calculated to assess reproducibility. One sample t-test was performed to evaluate changes induced by hypercapnia. We employed multiple linear regression to evaluate the association between age and CSF fraction in cortical gray matter, while adjusting for the effect of sex. The dependent variable was CSF fraction. Independent variables included age (years) and sex (male = 0, female = 1). Age was treated as a continuous predictor, and sex was coded as a binary categorical variable. All analyses were conducted in MATLAB using fitlm function. Statistical significance was defined as p < 0.05 (two-tailed).

## Supporting information

Extendeddatafigures

## Acknowledgements

This work was supported by the National Institutes of Health under award numbers U01 NS100588, R01 AG085299, R01 AG071515, R01 AG091312, P41 EB031771, U24 NS141774 and P30 AG066512.

## Supplementary information

Extendeddatafigures.docx

