## Extendeddatafigures for "Non-invasive MRI mapping of tissue-CSF water exchange reveals glymphatic fluid movement in live human cortex"

**Extended data figures**

**Extended Data Fig. 1: T2 decay characteristics of tissue and CSF compartments.**

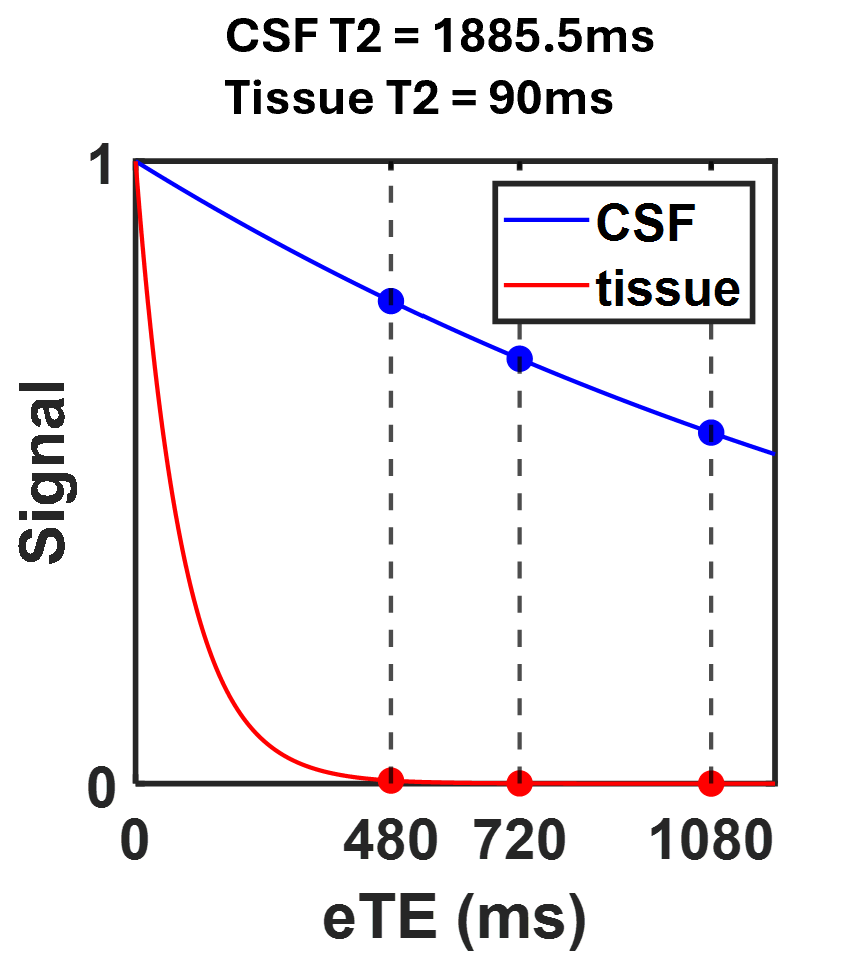

Simulation of signal attenuation as a function of effective echo time (eTE) for tissue (T2 = 90ms) and CSF (T2 = 1885.5ms). At the selected experimental eTE values (480, 720, and 1080ms), the tissue signal is predicted to decay to 0.483%, 0.034%, and 0.001% of the eTE = 0ms baseline, respectively. In contrast, the CSF signal remains largely preserved at 77.5%, 68.3%, and 56.4% of its initial magnitude. This differential decay allows for the selective isolation of ASL-labeled spins that have exchanged into the perivascular CSF space.

**Extended Data Fig. 2: Comparison of spatial patterns between perfusion and exchange maps.**

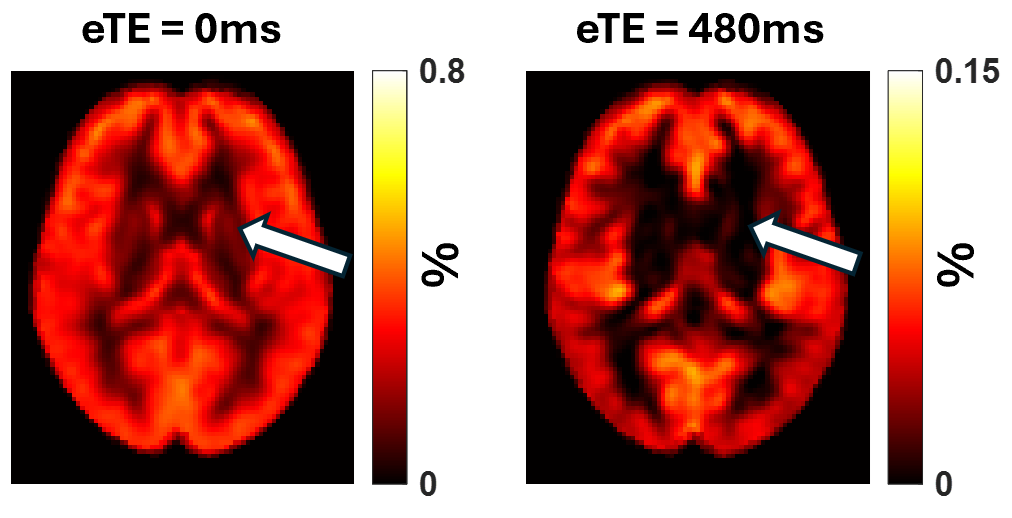

Visual comparison of the eTE = 0ms ASL map (brain perfusion) and the eTE = 480ms ASL map (perivascular exchange). White arrows highlight that while perfusion signals are prominent in deep gray matter structures such as the caudate, putamen, and thalamus, the long-TE ASL signal is markedly diminished in these regions compared to cortical gray matter.

**Extended Data Fig. 3: Anatomical definitions for regional analysis.**

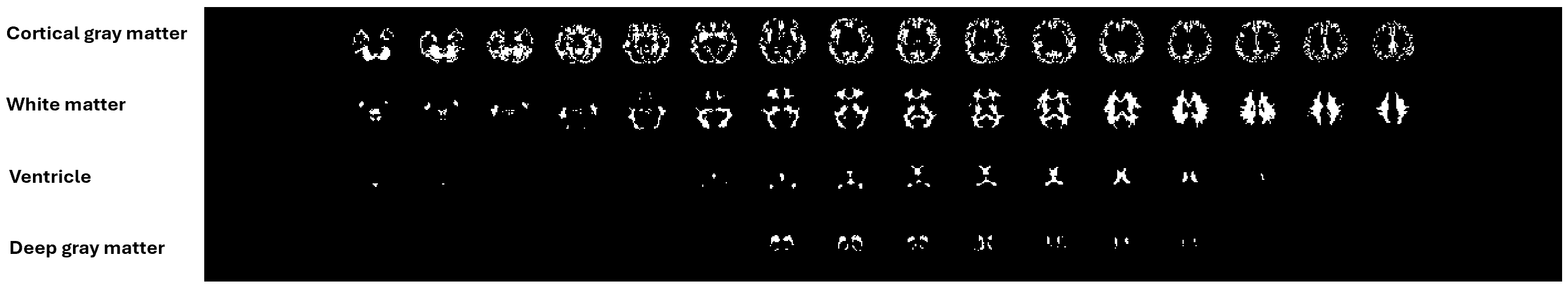

Four primary regions of interest (ROIs): cortical gray matter, white matter, deep gray matter (including caudate, putamen, and thalamus), and ventricles. These ROIs were derived semi-automatically for each subject from tissue probability maps generated from the eTE = 0ms M0 image using SPM12. Final definitions were established by thresholding the probability maps in combination with manual delineation to ensure anatomical accuracy.
